# Learning Minimal Gene Programs for Disease-Aligned Representations

**DOI:** 10.64898/2026.07.17.739212

**Authors:** Adithya V. Madduri, Chirag J. Patel

## Abstract

Identifying small, interpretable gene sets that robustly capture disease-associated variation in singlecell transcriptomic data remains a central challenge for biological interpretation and experimental followup. In practice, commonly used differential expression and sparsity-based approaches often produce large, unstable gene lists that fail to generalize across patients due to strong donor-specific confounding.

We study sparse gene selection for reconstructing donor-robust, disease-aligned cellular trajectories in real single-cell RNA-seq datasets. We introduce Sparse Linear Manifold Control (SLMC), a practical workflow that defines a disease-aligned score after removing donor-associated variation and selects minimal gene programs whose expression reconstructs this score. We focus on diagnosing the structure of the resulting reconstruction objective and evaluating selection strategies under realistic health data conditions.

Across five human single-cell datasets spanning oncology and neurodegeneration, we find that the reconstruction objective exhibits strong diminishing returns, explaining why simple greedy selection methods perform well in practice. Under strict donor-heldout evaluation, greedy methods consistently outperform LASSO at small gene budgets and achieve accurate reconstruction with as few as 25 genes. Together, these results highlight how careful objective design and empirical evaluation enable robust and interpretable gene selection for disease-aligned representation learning in single-cell health data.

**Data and Code Availability:** All code used for data preprocessing, model training, evaluation, and feature importance analyses is available at: https://github.com/AdiVM/SLMC_single-cell. The singlecell data used in this study are publicly available human transcriptomic datasets generated by prior studies and accessible through the Gene Expression Omnibus (GEO). Analyses were performed using renal cell carcinoma single-cell RNA-seq data (GEO accession: `GSE314072`) and Alzheimer’s disease single-nucleus RNA-seq data from human cortex (GEO accession: `GSE138852`), with additional publicly available datasets used for cross-context diagnostic evaluation (GEO accessions: `GSM8652069`, `GSE308624`, and `GSE227734`). All datasets contain de-identified human samples and are available under standard public-use terms via GEO.

**Institutional Review Board (IRB):** This study analyzes de-identified, publicly available human transcriptomic data obtained from previously published studies. No new data were collected, and no identifiable private information was accessed. In accordance with institutional policy, this work was determined to constitute non-human subjects research and did not require additional IRB approval.

## 1 Introduction

Identifying interventions that can reverse or attenuate disease-associated cellular states is a central challenge in medicine. Many successful therapies ultimately act by perturbing a small number of molecular pathways, yet pinpointing which genes or regulators to target remains difficult in complex, heterogeneous human tissues. Recent advances in singlecell transcriptomics have made it possible to measure disease-associated changes at the resolution of individual cells, revealing fine-grained alterations in specific cell types and states that are invisible in bulk tissue averages [Mathys et al., 2019, Miyoshi et al., 2024]. These technologies have transformed our ability to characterize disease, but they do not by themselves solve the key translational problem: which minimal set of genes should be perturbed to move a diseased cell toward a healthy state?

Standard differential expression analyses identify large lists of disease-associated genes, often numbering in the hundreds or thousands [Squair et al., 2021]. While informative, such lists are poorly suited for actionable intervention design, where practical constraints demand compact and interpretable target sets for follow-up experiments, drug development, or combinatorial perturbation screens. Moreover, human single-cell datasets are strongly confounded by inter-individual (donor) variation, technical batch effects, and cell-type composition differences. As a result, genes that appear predictive within one cohort may fail to generalize to new patients, limiting their clinical utility [Hou et al., 2023].

These challenges motivate the need for methods that (i) extract a disease-relevant trajectory at singlecell resolution, (ii) explicitly remove donor-specific confounding so that signals generalize across patients, and (iii) identify minimal gene programs that faithfully reconstruct this disease trajectory. Such sparse, donor-robust programs could serve as compact perturbation targets, candidate biomarker panels, or inputs to downstream generative and interventional models, enabling patient-robust hypothesis generation for therapeutic intervention.

In this work, we formulate sparse gene selection as reconstruction of a donor-orthogonal, disease-aligned manifold in single-cell transcriptomic space. We introduce Sparse Linear Manifold Control (SLMC), which first defines a disease progression axis in a structured PCA embedding and then explicitly removes donor-associated subspaces to isolate variation that is shared across patients. This yields a scalar, donor-robust disease score for every cell. We then search for small gene sets whose expression alone can reconstruct this disease score, producing compact and interpretable gene programs aligned with disease biology but invariant to donor identity. Finding these small gene sets is inherently a combinatorial optimization problem, since different subsets of genes can produce similar reconstructions of the disease trajectory. Although selecting the optimal subset of genes is combinatorial and NP-hard in general, we find that the resulting reconstruction objective has a highly structured geometry in real single-cell data. Across five diverse human datasets spanning cancer, neurodegeneration, and immune and cardiac contexts, the objective exhibits strong diminishing returns, with rare and small violations of submodularity. This near-submodular structure implies that simple greedy algorithms can efficiently recover near-optimal sparse gene programs without exhaustive search.

Empirically, we show that greedy selection of as few as 25–50 genes, which are small enough to be directly testable in combinatorial perturbation experiments, is sufficient to reconstruct donor-robust disease trajectories in both renal cell carcinoma and Alzheimer’s disease single-cell datasets. Gene sets selected under a strict donor-held-out protocol generalize to unseen patients, preserving both the ordering and scale of the disease-aligned cellular axis. Moreover, these compact programs recover established disease genes and clinically relevant targets, demonstrating that sparsity and patient-level robustness need not come at the expense of biological fidelity.

In summary, this work makes three contributions. First, we introduce SLMC, a framework for constructing donor-orthogonal, disease-aligned cellular trajectories and selecting minimal gene programs that reconstruct them. Second, we provide extensive empirical evidence that the resulting reconstruction objective is approximately submodular across diverse human single-cell datasets, explaining why simple greedy algorithms are both efficient and effective. Third, we demonstrate that the resulting sparse gene programs generalize across donors and recover clinically meaningful disease genes, providing compact, interpretable targets for downstream perturbation and biomarker design. By thus isolating donor-robust, disease-aligned variation and compressing it into minimal gene programs, SLMC provides compact and interpretable targets that generalize across patients, enabling more reliable biomarker design and prioritization of candidate perturbations for therapeutic intervention.

## 2 Problem Formulation and Disease-Aligned Objective

We address the problem of sparse gene selection in single-cell transcriptomics. Our objective is to identify a minimal set of genes *S* whose expression captures the transition from a diseased state toward a healthy reference, effectively providing a shortlist for in silico perturbation screening. Let *X* ∈ ℝ^*n×p*^ denote a standardized gene expression matrix with *n* cells and *p* genes. Each cell *i* is associated with a disease label *c*_*i*_ ∈ {disease, healthy} and a donor/batch identifier *d*_*i*_.

### 2.1 Disease Direction in a PCA Embedding

We first project the data into a low-dimensional space:

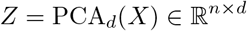

Let *I*_*D*_ and *I*_*H*_ denote the indices of diseased and healthy cells. We define the raw disease direction *v* as the vector connecting the centroids of these two populations in PCA space:

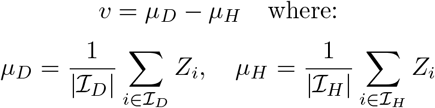

This vector *v* represents the primary axis of dysregulation. However, in multi-donor datasets, *v* is often confounded by technical batch effects or donor-specific biological variation.

### 2.2 Batch Artifact Correction and Donor Orthogonalization

To ensure our selection is clinically relevant and not driven by batch artifacts, we perform subspace pruning. We identify principal components (PCs) that represent “nuisance” variation. For each PC coordinate *Z*_*·k*_, we compute the fraction of variance explained (*R*^2^) by donor identity and disease status. PCs are treated as donor-confounded if they meet the following criteria:

- High donor association: *R*^2^(donor) *> τ*_donor_
- Low disease association: *R*^2^(disease) *< τ*_disease_

Both *R*^2^ quantities are computed directly from the data as the fraction of variance of each PC explained by the categorical donor or disease labels, so the identification of nuisance PCs is entirely data-driven rather than based on fixed component indices. Let *U* ∈ ℝ^*d×r*^ be the matrix whose columns are the *r* PCs satisfying these criteria. We define the donor-orthogonal disease direction *v*_*⊥*_ by projecting *v* onto the orthogonal complement of the donor subspace:

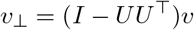

This operation selectively removes only those principal component directions that are empirically dominated by donor effects and weakly associated with disease, preserving disease-relevant variation while suppressing donor-specific structure. The thresholds *τ*_donor_ and *τ*_disease_ are fixed hyperparameters, but the set of removed components depends on the dataset-specific variance decomposition of each PC.

Finally, we assign each cell a scalar disease-aligned target score 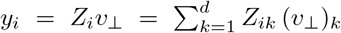, where the sum runs over the *d* principal-component coordinates of the embedding *Z*; this represents the “disease-ness” of a cell after removing donor-specific confounders.

### 2.3 Sparse Reconstruction Objective

We formulate gene selection as a combinatorial optimization problem. For a subset of genes *S* ⊆ {1, …, *p*}, we let *X*_*S*_ denote the corresponding columns of *X*. We evaluate *S* by its ability to reconstruct the target score *y* via ridge regression:

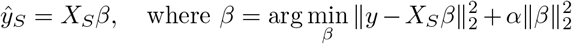

Our objective function *f* (*S*) is defined as the negative Mean Squared Error (MSE):

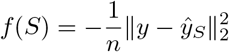

The goal is to find max_|*S*|*≤k*_ *f* (*S*). While this is NP-hard, the efficiency of greedy approximations depends heavily on the submodularity of *f* (*S*).

**Figure 1:**
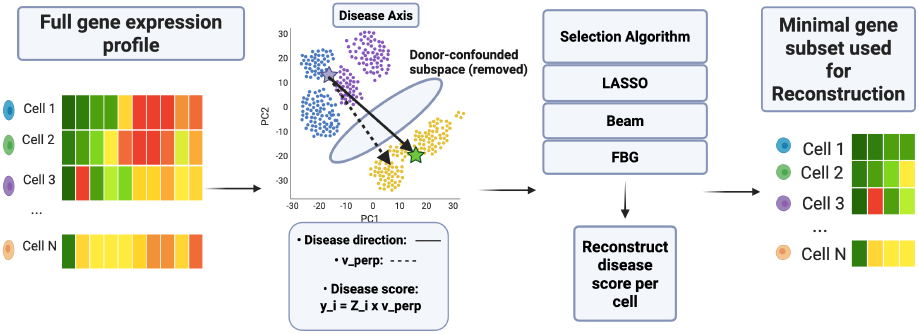
We define a donor-orthogonal disease axis in PCA space and project cells onto this axis to obtain a scalar disease score. Sparse gene selection methods are then used to identify small gene sets that reconstruct the disease score, yielding compact and interpretable representations of disease-aligned variation.

## 3 Related Work

### 3.1 Modeling and Predicting Cellular Perturbations

Recent deep generative models such as scGen enable prediction of whole-transcriptome responses to specified perturbations from single-cell data, allowing in silico simulation of cellular state transitions across cell types and conditions [Lotfollahi et al., 2019]. However, these approaches are optimized to accurately reconstruct the full gene expression response rather than to identify a minimal set of intervention targets. Compositional Perturbation Autoencoders extend this framework to model combinatorial interventions and covariates, but similarly operate in dense latent spaces and do not directly yield small, sparse gene panels tailored for experimental perturbation [Lotfollahi et al., 2021]. Gene regulatory network–based approaches such as CellOracle infer transcription factor–target interactions from single-cell multi-omics data and enable in silico simulation of cell identity changes under specified transcription factor perturbations, providing mechanistic insights into developmental and differentiation programs [Kamimoto et al., 2023]. These approaches focus on predicting the cellular consequences of specified perturbations, but do not directly address the inverse problem of identifying a small, donor-robust set of genes whose joint perturbation best drives cells along a disease-relevant trajectory.

### 3.2 Sparse and Interpretable Feature Selection in High Dimensions

Differential expression (DE) analysis in single-cell data has revealed cell-type–specific disease signals, although in most cases it identifies hundreds to thousands of disease-associated genes per cell type especially in large-scale human atlases [Mathys et al., 2019]. Subsequent work has also shown that careful handling of biological replicates is essential to avoid inflated false discoveries, as methods that ignore between-donor variation can report extensive DE even in the absence of true biological differences [Squair et al., 2021]. Even with well-calibrated testing procedures, trajectory-aware methods such as PseudotimeDE demonstrate that complex cellular progressions are typically associated with long, highly redundant gene lists rather than compact target sets [Song and Li, 2021]. In parallel areas of genomics, submodular optimization has been used to select small, diverse panels of assays that capture most of the relevant variation under strict budget constraints [Wei et al., 2016]. In contrast to DE-based ranking or assay panel selection, our approach identifies small, donor-robust sets of genes whose joint expression reconstructs disease-aligned cellular trajectories, yielding compact and experimentally actionable candidate perturbation targets in human disease settings.

### 3.3 Disease-Relevant Single-Cell Manifolds and Robust Generalization

Large-scale spatial and single-nucleus atlases of human disease demonstrate that transcriptomic alterations follow reproducible, structured progressions across tissues, regions, and donors, indicating the existence of shared disease-aligned cellular trajectories rather than donor-specific idiosyncrasies [Miyoshi et al., 2024]. Methodologically, recent statistical frameworks for multi-sample pseudotime analysis show that such trajectories must explicitly account for between-donor variability to avoid sample-specific effects that fail to generalize [Hou et al., 2023]. In parallel, deep integration models learn batch-invariant latent representations that align cells across datasets while preserving biological structure, enabling joint analysis in a shared manifold [Danino et al., 2024]. Complementarily, gene-level trajectory alignment methods reveal where dynamic programs agree or diverge between conditions, providing fine-grained comparisons of cellular progressions across systems [Sumanaweera et al., 2025]. Together, these lines of work establish that biologically meaningful and cross-donor comparable single-cell trajectories can be constructed and analyzed. Building on this foundation, our goal is to identify minimal gene programs whose expression alone reconstructs a donor-robust, disease-aligned trajectory that generalizes to unseen patients.

## 4 Methods

### 4.1 Empirical Submodularity Diagnostics

A set function is submodular if it exhibits “diminishing returns”: adding a gene to a small set provides more value than adding it to a larger set. To verify this property, we perform randomized triplet tests of the inequality:

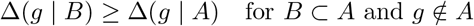

where Δ(*g* |*S*) = *f* (*S ∪* {*g*}) − *f* (*S*) is the marginal gain of adding gene *g*.

#### External-Manifold Diagnostic

To establish a baseline for single-cell manifolds, we split genes into “target” and “feature” sets. We reconstruct the PCA manifold of the target genes using the feature genes. This determines if the ridge reconstruction objective is inherently submodular in transcriptomic data.

#### Disease-Aligned Diagnostic

We repeat the test using the specific target *y* (from Section 2.2), for the two datasets used in our experiment. We report the Violation Rate (frequency where Δ_*B*_ *<* Δ_*A*_) and Violation Magnitude.

### 4.2 Datasets

We analyzed a total of five publicly available human single-cell datasets to assess the behavior of our external-manifold diagnostics across heterogeneous biological contexts. These datasets included: (i) GSM8652069, a healthy peripheral blood mononuclear cell (PBMC) single-cell multi-omic dataset from human blood; (ii) GSE314072, a renal cell carcinoma (RCC) single-cell study profiling tumor-infiltrating T cells; (iii) GSE308624, a spatial singlecell transcriptomics dataset of H. pylori–associated gastric cancer [Chen et al., 2025]; (iv) GSE227734, a single-cell cardiomyocyte dataset investigating hypertrophic cardiomyopathy (HCM) [Chen et al., 2024]; and (v) GSE138852, a single-nucleus RNA-seq atlas of the human cortex in Alzheimer’s disease [Grubman et al., 2019]. We selected two of these datasets, GSE314072 (RCC single-cell RNA-seq) and GSE138852 (Alzheimer’s disease single-cell RNA-seq) for analysis and validation using the experimental protocol outlined in 4.5. The RCC dataset (GSE314072) comprises 105,553 single cells, including 59,982 RCC cells and 45,571 healthy cells, while the Alzheimer’s disease dataset (GSE138852) consists of 13,214 nuclei, including 6,673 AD and 6,541 control nuclei. We selected GSE314072 and GSE138852 for donor-held-out generalization experiments.

### 4.3 Sparse Gene Selection Algorithms

Motivated by the empirical evidence of near-submodularity (violation rates *<* 10%), we compare four sparse gene selection strategies:

- **Forward Greedy (OMP-style):** Iteratively adds the gene *g* that maximizes the marginal gain Δ(*g* | *S*) = *f* (*S ∪* {*g*}) − *f* (*S*). This procedure is analogous to orthogonal matching pursuit in linear models, applied here to ridge-based reconstruction of the disease-aligned target.
- **Forward–Backward Greedy (FBG):** Extends forward greedy selection with a backward pruning step. After each addition, previously selected genes are removed if their removal does not decrease the objective beyond a small tolerance, yielding more parsimonious sets when redundancy arises.
- **Beam Search:** Maintains a fixed-width set of candidate solutions at each sparsity level, expanding multiple hypotheses in parallel. This method explores potential non-greedy interactions and serves as a stress test for the greedy assumption.

### 4.4 LASSO baseline (continuous *ℓ*_1_ relaxation)

As a continuous sparsity baseline, we evaluate LASSO regression, which replaces the combinatorial subset constraint with an *ℓ*_1_ penalty [Tibshirani, 1996]. Specifically, we solve

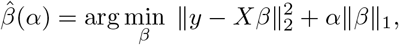

where *X* denotes the standardized gene expression matrix and *y* is the disease-aligned target. All genes are standardized to zero mean and unit variance prior to fitting.

We sweep the regularization parameter *α* over a logarithmically spaced grid of 50 values, ranging from the smallest value that yields a nonzero solution to a value that sets all coefficients to zero. Each *α* induces a model with a variable number of nonzero coefficients, corresponding to an implicit gene set size *k*(*α*).

To enable fair comparison with combinatorial methods at fixed gene budgets *k* ∈ {10, 25, 50, 100}, we select, for each target budget *k*, the LASSO model with *k*(*α*) *≤ k* that achieves the lowest reconstruction error. No refitting is performed after feature selection; coefficients are taken directly from the LASSO solution. Performance is evaluated using the same reconstruction metrics (MSE and *R*^2^) as for greedy and beam search methods.

### 4.5 Experimental Protocol

We evaluated all methods across five diverse singlecell datasets. For each dataset, we follow a consistent experimental protocol:

- **Sparsity sweep:** We evaluate gene set sizes *k* ∈ {10, 25, 50, 100}.
- **Robustness:** Each experiment is repeated across five random seeds, controlling randomness in PCA initialization, gene splits, and submodularity diagnostics.
- **Metrics:** We report mean squared error (MSE), which corresponds directly to the optimized objective, as well as the coefficient of determination (*R*^2^).

### 4.6 Donor-held-out generalization protocol

To evaluate whether sparse gene sets selected under the SLMC objective generalize beyond donor-specific structure, we adopt a donor-held-out evaluation scheme. For each dataset and random seed, donors are randomly partitioned into disjoint training (70%) and test (30%) sets, and all cells from a given donor are assigned exclusively to one split.

All preprocessing steps that could induce information leakage—including gene standardization, PCA embedding, donor-subspace identification, and construction of the disease-aligned target—are performed using training donors only. The learned PCA transformation and donor-orthogonal disease direction are then applied to held-out donors to compute test targets.

Sparse gene selection is performed exclusively on the training split. Generalization performance is assessed by evaluating the ridge reconstruction error of the learned gene sets on held-out donors. This protocol ensures that reported test performance reflects true out-of-donor generalization rather than memorization of donor-specific effects.

### 4.7 GWAS overlap analysis

To provide an external biological reference for the gene programs identified by each method, we assessed overlap with disease-associated genes reported in genome-wide association studies (GWAS). GWAS associations were obtained from the NHGRI–EBI GWAS Catalog (https://www.ebi.ac.uk/gwas/) for Alzheimer’s disease and renal cell carcinoma. For each disease, we extracted all genes listed in the `MAPPED GENE` field of the catalog associations, parsing comma-separated entries into individual gene symbols. For each method and dataset, we constructed a gene set by taking the union of genes selected across five random seeds at sparsity level *k* = 50. Overlap was computed as the set intersection between the selected gene set and the corresponding disease-specific GWAS gene set. This analysis was performed post hoc and was not used to guide gene selection or model optimization.

## 5 Results

### 5.1 Near-submodularity of single-cell gene reconstruction objectives

We evaluated whether the ridge reconstruction objective used for gene selection exhibits diminishing returns across five single-cell RNA-seq datasets. Using 2,000 randomized triplet tests per seed and five random seeds per dataset, mean violation rates ranged from 2.12% to 9.06% with low variance across seeds (standard deviation *≤* 1.44%; Table 1). In all datasets, the mean marginal difference Δ(*g* |*S*^*′*^) − Δ(*g* | *S*) was negative. We did observe rare large violations in two datasets (GSE227734 and GSM8652069); however, these events were directly attributable to numerically ill-conditioned ridge fits arising from extreme gene–sample imbalance, occurred with negligible frequency, and did not affect aggregate violation rates. Overall, these results indicate that the reconstruction objective is approximately submodular, supporting greedy and forward–backward selection.

**Table 1:** Submodularity diagnostics across datasets.

| Dataset | Violation rate (%) |
| --- | --- |
| GSE138852 | 5.75 |
| GSE227734 | 5.22 |
| GSE308624 | 3.62 |
| GSE314072 | 2.12 |
| GSM8652069 | 9.06 |

#### 5.2 Predictive performance of FBG versus baselines

We compared Forward–Backward Greedy (FBG), pure Greedy, Beam Search, and LASSO on two representative datasets (GSE314072 and GSE138852) across gene budgets *k* ∈ {10, 25, 50, 100} and five random seeds.

On the RCC dataset (GSE314072), FBG, Greedy, and Beam exhibited nearly identical performance across all budgets. At *k* = 10, all three methods achieved the same performance (MSE 268.866, *R*^2^ 0.859), while LASSO performed substantially worse (MSE 685.213, *R*^2^ 0.640). At *k* = 50, FBG achieved an MSE of 99.804 compared to 100.657 for Beam, with identical *R*^2^ of 0.947. At *k* = 100, FBG achieved an MSE of 57.574 and *R*^2^ of 0.970, marginally improving over Beam (MSE 59.404).

On AD dataset (GSE138852), we observed patterns highly consistent with the RCC dataset. At *k* = 25, FBG achieved a reconstruction MSE of 113.661 (*R*^2^ = 0.916), outperforming Beam Search (MSE 115.588). This performance gap persisted at higher gene budgets; by *k* = 100, FBG reached an MSE of 43.105 (*R*^2^ = 0.968), while Beam Search trailed at 48.813. Across all sparsity levels, the continuous LASSO baseline underperformed significantly, yielding MSE values 2–5 *×* larger than combinatorial methods at low *k*.

Taken together, these results indicate that forward greedy selection is sufficient to attain near-optimal predictive performance under the reconstruction objective studied here. FBG and Beam search do not provide systematic improvements over pure Greedy, but closely track its performance, consistent with an objective that exhibits strong diminishing returns. In contrast, convex sparsity-based selection via LASSO performs substantially worse at small gene budgets.

### 5.3 Marginal gain decay and diminishing returns during selection

We analyzed the trajectory of marginal gains during FBG selection on GSE314072 at *k* = 50. The mean marginal gain at step 1 was 1.03 *×* 10^3^, dropping to 2.68 *×* 10^2^ by step 2 and 1.11 *×* 10^2^ by step 3. By step 10, the marginal gain had decreased to 1.62 *×* 10^1^.

When normalized by the first step, the relative marginal gain dropped to 0.26 at step 2, 0.11 at step 3, and 0.016 by step 10. At the midpoint (step 25), the relative gain was 0.004, and by the final step it was 0.001. Across 49 transitions, only 5 monotonicity violations were observed, confirming near-monotonic decay. This quantitative decay strongly supports the presence of diminishing returns consistent with approximate submodularity.

### 5.4 Out-of-donor generalization testing

We evaluated out-of-donor generalization of sparse gene sets selected under the SLMC objective using donor-held-out evaluation on two biologically distinct datasets: RCC (GSE314072) and Alzheimer’s disease (AD; GSE138852). Across five random donor splits and gene budgets *k* ∈ {10, 25, 50, 100}, test performance increased monotonically with *k* for all methods, indicating that larger gene sets consistently improve reconstruction of the disease-aligned axis under donor hold-out.

**Figure 2:**
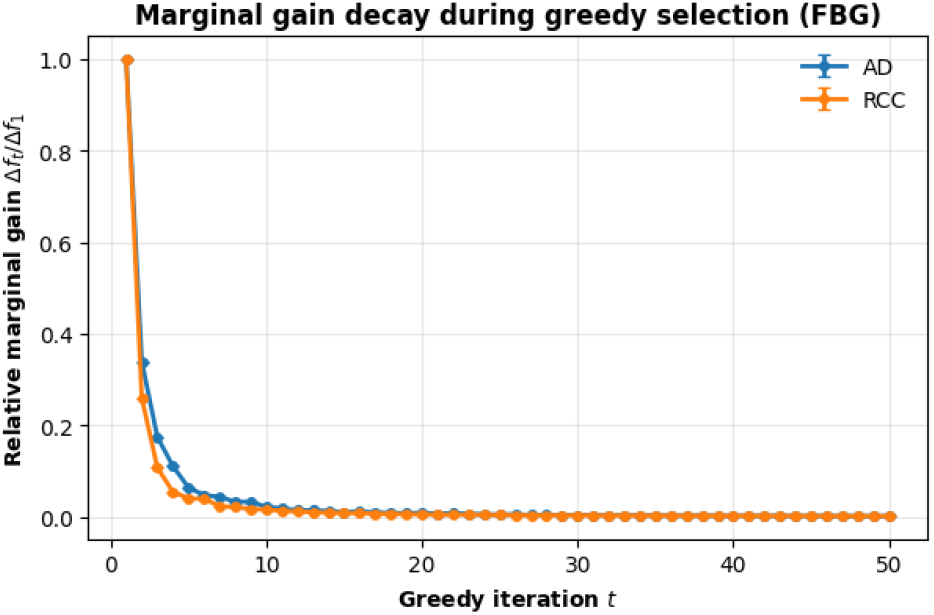
Marginal gain decay during forward– backward greedy (FBG) selection at *k* = 50. Relative marginal gains decrease rapidly across greedy iterations, with near-monotonic decay across seeds and datasets, indicating strong diminishing returns consistent with approximate submodularity.

In RCC, Forward–Backward Greedy (FBG) closely matched beam search across all sparsity levels, while substantially outperforming LASSO under tight sparsity. In AD, overall generalization was weaker and variance across donor splits was higher, reflecting greater biological heterogeneity; nevertheless, FBG again outperformed LASSO at all budgets and consistently exceeded beam search at small and intermediate *k*, with all methods converging at larger *k*. These results show that greedy optimization under the SLMC objective yields sparse gene sets that generalize robustly across donors while preserving the geometry of the disease-aligned trajectory.

**Figure 3:**
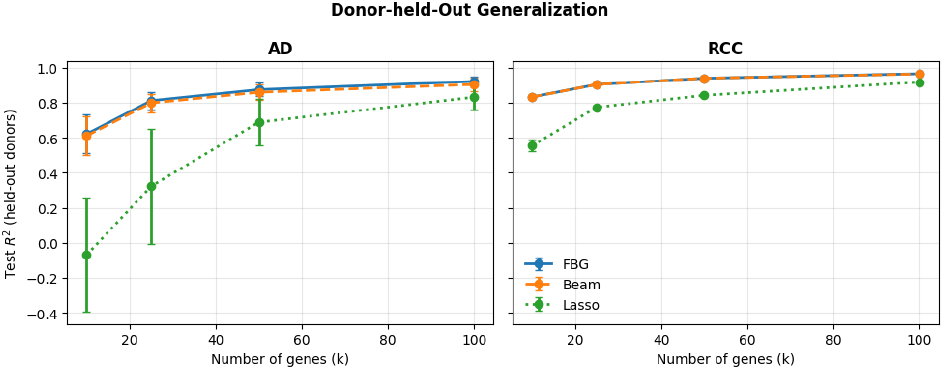
Out-of-donor generalization under the SLMC objective. Test *R*^2^ on held-out donors is shown as a function of gene budget *k* for FBG, Beam Search, and LASSO, averaged across five donor splits, for RCC and AD datasets.

**Figure 4:**
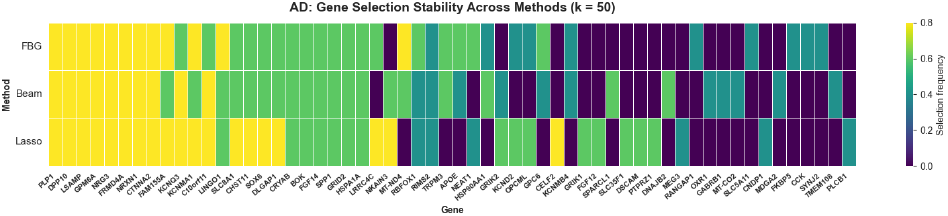
Gene selection stability across methods in Alzheimer’s disease at *k* = 50. Each column corresponds to a gene, and color intensity indicates selection frequency across five donor-held-out splits.

### 5.5 Biological and Clinical Validation of Selected Gene Programs

To assess the biological relevance of the identified gene programs, we evaluated the overlap between genes selected across all five random donorheld-out splits (*k* = 50) and curated sets of disease-associated genes from Genome-Wide Association Studies (GWAS) [Sollis et al., 2023]. In the Alzheimer’s Disease (AD) context, Forward-Backward Greedy (FBG) identified 26 GWAS-associated genes, achieving parity with the more computationally intensive Beam Search (26 genes) and outperforming LASSO (23 genes). Notably, FBG consistently prioritized “gold standard” AD risk genes, including **APOE, MEF2C, NRXN1**, and **RBFOX1** [Ren et al., 2022, Bellenguez et al., 2022]. High-confidence drivers such as **LINGO1** and **APOE** exhibited higher selection frequencies across donor-held-out splits in the FBG model compared to both Beam Search and LASSO [Fernandez-Enright and Andrews, 2016, Jackson et al., 2024]. This demonstrates that the approximately submodular geometry of the SLMC objective allows greedy trajectories to capture the core genetic architecture of the disease manifold as effectively as complex procedures, while remaining invariant to inter-donor variation.

In the Renal Cell Carcinoma (RCC) context, while the current GWAS catalog remains relatively underdeveloped for this disease—yielding zero overlaps for all models at *k* = 50—the gene programs identified by FBG are highly enriched for emerging therapeutic targets and biomarkers. Specifically, FBG identified **LAG3**, a primary immune checkpoint protein currently the subject of Phase II clinical trials for dual blockade with PD-1 in RCC patients [Qiu et al., 2024]. Other identified genes include **TNFRSF9**, a co-stimulatory receptor on CD8+ T cells, and **UBE2C**, a member of the ubiquitin modification system recently shown to promote tumor cell proliferation and predict poor overall survival in RCC [Chen and Wang, 2021]. Additionally, FBG prioritized **CLIC1**, which has been correlated with tumor grade, metastasis, and poor prognosis in clear-cell RCC [Nesiu et al., 2019]. The robust identification of these clinically significant candidates under strict donor-held-out generalization testing underscores the utility of the SLMC framework for prioritizing interpretable gene sets that reflect actionable biological trajectories.

**Figure 5:**
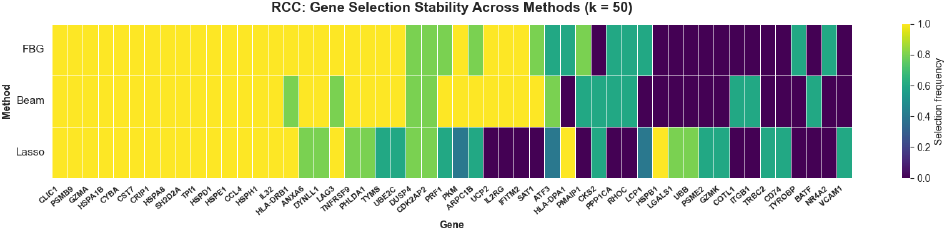
Gene selection stability across methods in RCC at *k* = 50. Each column corresponds to a gene, and color intensity indicates selection frequency across five donor-held-out splits.

## 6 Discussion

This work studies a practical but underexplored problem in single-cell transcriptomics: identifying small, donor-robust gene programs that faithfully reconstruct disease-aligned variation rather than merely ranking genes by marginal association. Our results show that, across multiple human datasets, a simple linear reconstruction objective defined on a donor-orthogonal disease axis admits highly sparse solutions that generalize to unseen donors. Empirically, we find that as few as 25–50 genes are sufficient to recover the ordering and scale of disease-aligned cellular trajectories in both renal cell carcinoma and Alzheimer’s disease. These findings suggest that compact gene programs can capture meaningful disease structure without requiring dense latent representations or complex nonlinear models, provided that donor-specific confounding is explicitly addressed.

A central contribution of this work is the observation that the SLMC reconstruction objective exhibits strong diminishing returns in real single-cell data. Although greedy feature selection for regression is not submodular in general, we empirically demonstrate low violation rates and small violation magnitudes across five heterogeneous datasets, consistent with weak submodularity under ridge regularization. This structure explains why simple forward greedy selection performs comparably to more computationally intensive beam search, and why backward pruning is rarely triggered in practice. These findings complement prior theoretical results on weak submodularity in sparse regression objectives and help bridge the gap between approximation guarantees and observed behavior in high-dimensional biological data [Wei et al., 2016].

From an applied perspective, our evaluation protocol emphasizes robustness and generalization rather than in-sample fit. By adopting strict donor-heldout evaluation, we show that sparse gene programs selected under SLMC generalize beyond individual-specific effects, a known challenge in human single-cell studies [Squair et al., 2021]. In both RCC and AD, greedy-selected gene sets preserve disease-aligned structure on unseen donors, while continuous *ℓ*_1_ relaxation via LASSO underperforms at small gene budgets. Importantly, the biological validation via GWAS overlap and literature-supported targets demonstrates that donor robustness and interpretability need not come at the expense of disease relevance, particularly when selection is constrained to small, experimentally actionable gene sets.

More broadly, this work aligns with a growing body of research emphasizing careful evaluation and robustness in machine learning for health applications. Rather than proposing a new predictive architecture, we focus on diagnosing objective structure, benchmarking selection strategies, and stress-testing generalization under realistic confounding. This emphasis on evaluation, interpretability, and robustness situates SLMC as a practical tool for hypothesis generation and downstream experimental prioritization, complementing existing approaches that model dense transcriptomic responses or simulate perturbations without explicitly enforcing sparsity [Lotfollahi et al., 2019, 2021, Kamimoto et al., 2023].

Ultimately, the value of identifying donor-robust, disease-aligned gene programs lies in their potential to inform intervention. By isolating small sets of genes whose joint expression captures disease-associated cellular states across patients, our approach yields compact candidate targets that are directly compatible with downstream experimental and clinical workflows.

### 6.1 Limitations

Our study has several important limitations. First, SLMC relies on a linear reconstruction objective defined in a PCA embedding, which may fail to capture nonlinear gene interactions or branching disease trajectories present in some biological systems. In practice, such nonlinear or branching structure would manifest as poor reconstruction of the disease-aligned score and rapidly saturating performance with increasing gene budgets, signaling that SLMC’s linear assumptions are insufficient for the dataset at hand. Second, the choice of PCA as the embedding space, while effective for denoising and capturing global structure, may discard biologically meaningful variation that resides in lower-variance components. Third, our GWAS overlap analysis is descriptive rather than inferential; overlap should be interpreted as contextual biological support rather than validation of causal relevance. Furthermore, in highly heterogeneous settings such as Alzheimer’s disease, we observe reduced stability of very small gene sets, indicating that multiple distinct but similarly predictive programs may exist and that extreme sparsity may amplify sensitivity to donor composition. Additionally, in this work we do not explicitly model cell-type–specific trajectories. While such stratification may be important in highly heterogeneous diseases such as Alzheimer’s disease, applying SLMC within individual cell types would substantially reduce effective sample sizes and stability under tight sparsity constraints; in contrast, for settings such as renal cell carcinoma, where the data are dominated by a small number of immune cell populations, cell-type–agnostic modeling is less restrictive and biologically appropriate.

Our analyses are primarily descriptive rather than inferential. The reported PC-wise donor and disease *R*^2^ values, submodularity violation rates and magnitudes, held-out *R*^2^/MSE differences, gene-selection frequencies, and empirical RSC/RSM ratios are point estimates or summaries across a small number of data splits. Because cells are nested within donors, and donors may recur across splits, neither cells nor random seeds constitute independent biological replicates. Consequently, the reported variability does not provide uncertainty for claims concerning donor orthogonality, near-submodularity, gene-set stability, or comparative predictive performance. Future analyses should use donor-clustered bootstrap or permutation procedures, paired donor-level comparisons, or hierarchical models, together with sensitivity analyses over PCA dimensionality and nuisance-PC thresholds.

### 6.2 Future Directions

Several extensions of our work are promising. While SLMC relies on linear reconstruction objectives for interpretability and stability, incorporating nonlinear or kernelized variants could capture higher-order gene interactions while retaining explicit sparsity constraints, potentially improving fidelity when disease progression is not well approximated by a single linear axis. Extending SLMC to cell-type–specific trajectory modeling is another natural direction: restricting analysis to individual cell types may yield more mechanistically specific gene programs, particularly in heterogeneous diseases such as Alzheimer’s disease, but must be balanced against reduced sample sizes and increased variance that can limit stability under tight sparsity constraints. The recovery of canonical disease-associated genes in the present cell-type–agnostic setting suggests that explicit stratification is not strictly required, but could provide additional resolution when sufficient data are available. Beyond transcriptomic reconstruction, coupling SLMC-selected gene programs with perturbation-response models or CRISPR screening data could enable direct evaluation of causal effects along disease-aligned axes [Dixit et al., 2016], while extension to longitudinal or multi-modal datasets (e.g., proteomic or clinical data) offers a path toward aligning compact, donor-robust programs with disease progression over time rather than static case–control contrasts.

## 7 Conclusion

Our work addresses a critical bottleneck in precision medicine: the gap between high-dimensional, donor-confounded single-cell disease signals and the small, stable gene programs required for robust interpretation, evaluation, and downstream experimental use. We study how disease-aligned objectives behave in practice under realistic confounding and which sparse selection strategies reliably recover patient-robust signal.

Through systematic out-of-donor evaluation across multiple human single-cell datasets, strict donorheld-out protocols, and explicit diagnostics of objective structure, we show that simple greedy selection is sufficient to identify minimal gene programs that generalize across patients and preserve disease-aligned trajectories. These results clarify why commonly used continuous sparsity relaxations underperform at small budgets, and provide concrete guidance for practitioners selecting compact, interpretable gene sets in applied health settings.

By emphasizing evaluation, robustness, and interpretability, our findings position sparse gene program selection as a practical diagnostic and benchmarking task for single-cell disease analysis. We expect this perspective to support more reliable comparison of methods, clearer experimental prioritization, and closer alignment between machine learning analyses and downstream precision medicine workflows.

## A. First Appendix

### A.1 Theoretical Justification of Near Submodularity

Greedy subset selection for linear reconstruction objectives is not submodular in general; however, such objectives are known to satisfy weak submodularity under standard restricted eigenvalue conditions on the design matrix [Das and Kempe, 2011, Elenberg et al., 2018]. Weak submodularity is quantified via the submodularity ratio, which measures the extent to which a set function exhibits diminishing returns. For objectives arising from (regularized) least-squares regression, prior work shows that the submodularity ratio is lower bounded by quantities depending on the restricted strong convexity (RSC) and restricted strong smoothness (RSS) constants of the objective over sparse supports [Elenberg et al., 2018, Khanna et al., 2017]. Concretely, if the underlying ridge regression loss is *m*-restricted strongly convex and *M* - restricted strongly smooth in the coefficients on supports of size at most *k*, then the induced set function *f* (*S*) has submodularity ratio bounded below by *m/M* [Elenberg et al., 2018, Khanna et al., 2017]. These conditions are strictly weaker than exact sub-modularity and hold for a broad class of design matrices encountered in practice, provided that the covariance structure is well-conditioned on sparse subsets.

In SLMC, the ridge-regularized least-squares loss

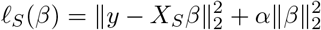

satisfies the RSC/RSS conditions whenever the regularized Gram matrix 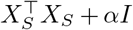 is uniformly well-conditioned over sparse supports [Elenberg et al., 2018, Khanna et al., 2017]. Under these conditions, the associated set function

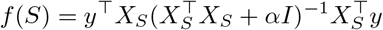

is weakly submodular [Elenberg et al., 2018, Khanna et al., 2017]. Ridge regularization therefore stabilizes the spectral properties governing the submodularity ratio by preventing arbitrarily small or large marginal gains induced by collinearity. Importantly, this theory does not imply that the SLMC objective is exactly submodular, nor does it guarantee that the submodularity ratio is close to one in the worst case. Rather, it establishes that greedy algorithms admit constant-factor approximation guarantees of the form *f* (*S*_greedy_) *≥* (1 − *e*^−*γ*^)*f* (*S*^*⋆*^), where *γ* is the submodularity ratio [Das and Kempe, 2011, Bian et al., 2017]. Our empirical results in Section 5 demonstrate that, for disease-aligned single-cell manifolds, observed violation rates and magnitudes are small, indicating that the effective submodularity ratio is high in practice—consistent with, but not implied by, the above theoretical bounds.

#### A.2 Complexity Analysis of Selection Algorithms

We compare the computational cost of SLMC-FBG to combinatorial search baselines. Let *n* denote the number of cells, *p* the number of candidate genes (capped at 5000 in our implementation), *k* the target sparsity level, and *B* the beam width.

- **Forward Greedy / FBG:** At each of the *k* selection steps, the algorithm evaluates all remaining *p* genes, resulting in *O*(*k · p*) ridge regression fits. Each ridge fit on a subset of size *s ≤ k* has cost *O*(*ns*^2^) (ignoring lower-order terms), yielding an overall complexity of *O*(*n · k*^3^ *· p*) in the worst case. Although FBG includes a backward pruning step, we empirically observe that backward removals are rarely triggered, making its runtime effectively identical to pure forward greedy selection in practice.
- **Beam Search:** Unrestricted beam search would explore *O*(*k · p · B*) candidates. To ensure tractability, our implementation restricts expansion to a candidate pool of size *M ≪ p* based on initial correlation with the target, reducing the cost to *O*(*k · M · B*) ridge fits. Despite this heuristic restriction, FBG—which evaluates the full feature space at each step—matches or exceeds Beam Search performance under the SLMC objective, indicating that greedy trajectories are sufficient in this near-submodular regime.
- **LASSO (continuous** *ℓ*_1_ **relaxation):** Our LASSO baseline performs a sweep over *A* regularization values (with *A* = 50 in our experiments), fitting a convex *ℓ*_1_-penalized regression model at each value. Each fit has cost approximately *O*(*n · p*) per coordinate descent pass, yielding a total complexity of *O*(*A · n · p*) for the sweep. While individual LASSO fits are computationally efficient, the method does not directly optimize for a fixed sparsity level *k* and requires post-hoc model selection to match target budgets, in contrast to the explicit *k*-controlled selection used by greedy and beam-based methods.

#### A.3 Out-of-donor generalization across datasets

We first evaluated whether sparse gene sets selected under the SLMC objective generalize to unseen donors using the RCC dataset (GSE314072). Across five random seeds and gene budgets *k* ∈ {10, 25, 50, 100}, combinatorial methods exhibited strong out-of-donor generalization, with test *R*^2^ increasing monotonically as a function of k. Forward–Backward Greedy (FBG) consistently matched beam search performance on held-out donors. Averaged across seeds, FBG achieved test *R*^2^ = 0.833 at k=10, 0.904 at k=25, 0.940 at k=50, and 0.966 at k=100, closely tracking beam search at all sparsity levels. In contrast, LASSO-based selection generalized substantially worse under tight sparsity constraints. At k=10, LASSO achieved mean test *R*^2^ = 0.55, compared to 0.83 for FBG. Although LASSO performance improved with increasing sparsity, it remained consistently below combinatorial methods across all budgets. This gap was most pronounced at small k, indicating that continuous *ℓ*_1_ relaxation underfits when the goal is to identify minimal gene sets that generalize across donors. Notably, the parity between FBG and beam search persisted under donor-held-out evaluation, despite the presence of mild submodularity violations measured on training data. This suggests that the near-submodular structure of the SLMC reconstruction objective reflects intrinsic geometry of the disease-aligned manifold rather than donor-specific artifacts. As a result, simple greedy selection is sufficient to identify gene sets that generalize across donors, without requiring more expensive combinatorial search. Together, these results demonstrate that the geometric structure underlying the SLMC objective not only supports efficient greedy optimization, but also yields sparse gene sets that generalize robustly across donors.

We next evaluated out-of-donor generalization of sparse gene sets under the SLMC objective using the Alzheimer’s disease dataset (GSE138852). As in the RCC setting, performance was assessed across five random donor-held-out splits and gene budgets *k* ∈ { 10, 25, 50, 100} . In contrast to RCC, overall generalization performance was lower and exhibited substantially higher variability across donor splits, reflecting the greater heterogeneity of AD samples.

For the combinatorial methods, test performance increased monotonically with increasing gene budget. Forward–Backward Greedy (FBG) achieved mean test *R*^2^ = 0.56 at *k* = 10, improving to 0.79 at *k* = 25, 0.85 at *k* = 50, and 0.91 at *k* = 100. Beam search followed a similar trend but consistently underperformed FBG at all sparsity levels, with mean test *R*^2^ = 0.55 at *k* = 10, 0.77 at *k* = 25, 0.84 at *k* = 50, and 0.88 at *k* = 100. Notably, both methods exhibited large standard deviations at small *k*, indicating sensitivity to donor composition under severe sparsity.

LASSO-based selection generalized poorly under tight sparsity constraints in the AD setting. At *k* = 10, LASSO achieved negative mean test *R*^2^ (− 0.25), indicating systematic underfitting on heldout donors. Performance improved with increasing sparsity, reaching mean test *R*^2^ = 0.13 at *k* = 25, 0.61 at *k* = 50, and 0.79 at *k* = 100, but remained substantially below combinatorial methods across all budgets. This pronounced gap at low *k* highlights the difficulty of recovering donor-robust signal in AD using continuous *ℓ*_1_ relaxation when constrained to minimal gene sets.

As in RCC, FBG required no backward deletions across all runs, while beam search exhibited no gain monotonicity violations by construction. Despite the presence of mild submodularity violations in the FBG forward gains (185 total across seeds and budgets), greedy selection consistently outperformed or matched beam search in terms of test performance. Together, these results indicate that although AD presents a more challenging and heterogeneous generalization setting than RCC, the SLMC objective retains sufficient near-submodular structure for simple greedy optimization to recover sparse gene sets that generalize across donors, particularly at moderate sparsity levels.

#### A.4 Held-out Donor Reconstruction Fidelity

To complement the aggregate donor-held-out generalization metrics reported in Section 5.4, we provide a qualitative and distribution-level assessment of reconstruction fidelity on unseen donors. Specifically, we visualize the true donor-orthogonal disease scores *y*_test_ against their sparse reconstructions *ŷ*_test_, pooling cells across five independent random donorheld-out splits (seeds) and multiple sparsity levels *k* ∈ {10, 25, 50, 100} . All quantities are computed using PCA embeddings, donor-subspace identification, and disease-aligned directions learned exclusively on training donors, and applied without refitting to heldout donors. Note that, unlike the split-averaged *R*^2^ reported above, reconstruction fidelity here is computed by pooling held-out cells across donor splits, yielding a single global *R*^2^ that reflects cell-level alignment rather than split-level variability.

Across both datasets, these visualizations confirm that high test *R*^2^ reflects faithful recovery of the disease-aligned axis itself—preserving both ordering and scale—rather than trivial rescaling or variance shrinkage. In the Alzheimer’s disease dataset, reconstruction fidelity increases monotonically with sparsity, achieving pooled held-out *R*^2^ values of 0.81, 0.91, 0.94, and 0.96 at *k* = 10, 25, 50, and 100, respectively, while maintaining strong disease discrimination (AUROC(*y*_test_) = 0.93; AUROC(*ŷ*_test_) = 0.86– 0.93). A similar pattern is observed in renal cell carcinoma, where held-out *R*^2^ increases from 0.85 at *k* = 10 to 0.97 at *k* = 100, with AUROC preserved between the true and reconstructed scores (AUROC(*y*_test_) = 0.91; AUROC(*ŷ*_test_) = 0.86–0.90).

Together, these results demonstrate that sparse gene sets selected under the SLMC objective not only generalize across donors in aggregate metrics, but also reconstruct the same donor-orthogonal disease trajectory at the level of individual cells, supporting the interpretability and validity of the donor-held-out generalization analysis.

**Figure 6:**
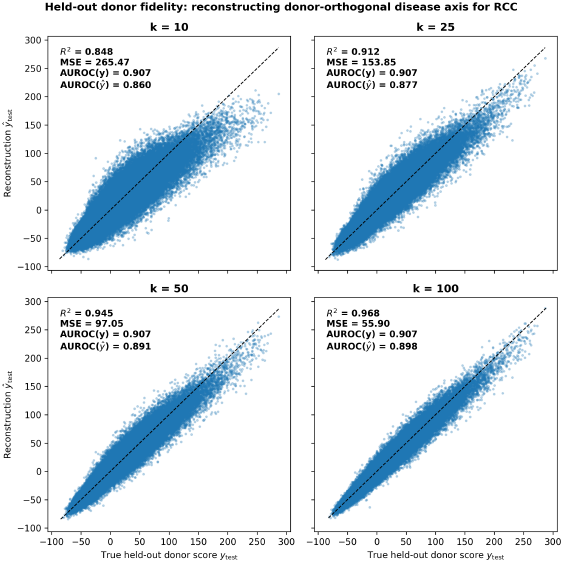
Held-out donor reconstruction fidelity for renal cell carcinoma (GSE314072). True donor-orthogonal disease scores *y*_test_ are plotted against sparse reconstructions *ŷ*_test_ for *k* ∈ {10, 25, 50, 100}, pooled across five donor-held-out splits.

#### A.5 Selection Stability via Jaccard Similarity

We evaluated selection stability using pairwise Jaccard similarity of the full selected gene sets across five random seeds under the out-of-donor protocol.

For the RCC dataset (GSE314072), stability was moderate and consistent across sparsity levels. At *k* = 10, the mean Jaccard similarity was 0.523 (std 0.118, range 0.333–0.667). Increasing the budget to *k* = 50 yielded a similar but slightly more concentrated overlap, with mean Jaccard similarity 0.545 (std 0.055, range 0.471–0.639).

In contrast, the AD dataset (GSE138852) exhibited lower and more variable stability. At *k* = 10, the mean Jaccard similarity was 0.382 (std 0.235, range 0.250–1.000), reflecting strong sensitivity of small gene sets to donor splits and initialization. At *k* = 50, the mean Jaccard similarity was 0.378 (std 0.194, range 0.250–0.887), again indicating limited but highly variable overlap between runs. The presence of occasional very high overlaps (e.g., Jaccard *≈* 0.89 or 1.00 for specific seed pairs) alongside many low-overlap pairs suggests that multiple distinct but similarly predictive gene programs can be recovered in the AD setting.

**Figure 7:**
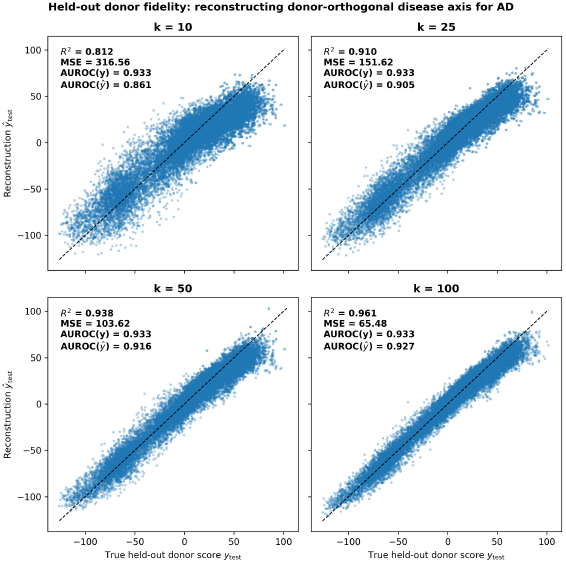
Held-out donor reconstruction fidelity for Alzheimer’s disease (GSE138852). True donor-orthogonal disease scores *y*_test_ are plotted against sparse reconstructions *ŷ*_test_ for *k* ∈ {10, 25, 50, 100}, pooled across five donor-held-out splits.

#### A.6 Optimization Diagnostics on Full-Dataset Fits

In this section we analyze the optimization behavior of FBG when fit on the full dataset without out-of-donor holdout. Unlike the main experiments, which evaluate generalization across unseen donors, these diagnostics are designed to probe intrinsic properties of the reconstruction objective and greedy search dynamics (e.g., stability, effective submodularity, and redundancy along the selection path). Evaluating on the full dataset removes cross-donor sampling noise and allows us to directly study how the objective behaves as genes are added or removed.

##### A.6.1 Role of backward steps in FBG

To assess the contribution of backward elimination in Forward–Backward Greedy (FBG), we explicitly tracked backward activity at both the step level and the run level. Across all random seeds and gene budgets (*k* ∈ { 10, 25, 50, 100}), no forward step triggered a backward removal, and no run exhibited any backward elimination events. As a result, the total number of genes removed per run was exactly zero in all experiments. Rather than indicating a failure of the backward mechanism, this behavior provides empirical evidence that the reconstruction objective behaves as effectively submodular in the regimes considered. In particular, once a gene is selected by the forward step, its marginal contribution is never outweighed by that of a later-added gene, making backward correction unnecessary. This observation is consistent with the low submodularity violation rates and small violation magnitudes observed in our empirical submodularity diagnostics.

Together, these results indicate that the greedy forward path is locally stable under the objective studied here: selections made by forward greedy are never reversed by backward pruning. Consequently, FBG closely matches pure forward greedy selection across both the RCC and AD datasets, not because backward steps are disabled, but because the objective itself renders them redundant.

##### A.6.2. Robustness to submodularity violations

We stratified runs by median violation rate and compared FBG and Beam at *k* = 50. In the low-violation regime, FBG achieved mean MSE 99.72 compared to 100.60 for Beam. In the high-violation regime, FBG achieved mean MSE 99.93 compared to 100.74 for Beam. This trend was even more pronounced in the AD dataset, where FBG maintained a *∼* 2.3 MSE lead over Beam search regardless of violation frequency. The performance gap remained consistent across regimes, indicating that FBG’s performance is robust to moderate deviations from exact submodularity.

##### A.6.3. Gene-level contribution structure and redundancy

We examined gene-level contributions along the FBG trajectory. Early selections yielded large marginal gains with moderate redundancy: mean relative gain over steps 1–10 was 0.158, with mean maximum correlation to previously selected genes of 0.390. In contrast, late selections (steps 41–50) exhibited mean relative gain of 0.002 and lower redundancy (mean correlation 0.316). Similar decay trajectories were observed in the AD context.

This indicates that early selections capture most of the predictive signal, while later additions contribute marginal refinements without excessive redundancy.

#### A.7 Robustness of Near-Submodularity Under Nonlinear Embeddings

To assess whether the empirical near-submodularity and out-of-donor generalization results reported in the main text are artifacts of the linear PCA embedding, we conducted two sets of experiments replacing linear PCA with a nonlinear RBF Kernel PCA embedding (50 components, fit on a subsample of 5,000 cells per training split for memory tractability). All other aspects of the pipeline were held exactly constant: the same donor splits (70*/*30 by patient), the same gene filtering and standardization, the same FBG selection algorithm, the same ridge regularization (*α* = 1.0), and the same five random seeds across gene budgets *k* ∈ {10, 25, 50, 100}.

##### Donor-Held-Out Generalization Under Kernel PCA

We reran the full donor-held-out generalization experiment on the RCC dataset (GSE314072) with Kernel PCA in place of linear PCA. Under linear PCA, FBG achieved mean test *R*^2^ of 0.833, 0.904, 0.940, and 0.966 at *k* = 10, 25, 50, 100, respectively (std *≤* 0.018 across seeds). Under RBF Kernel PCA, the corresponding mean test *R*^2^ values were 0.538, 0.593, 0.621, and 0.670 (std *≤* 0.132). The nonlinear embedding yields substantially lower out-of-donor generalization at every sparsity level, with a consistent gap of approximately 0.30 *R*^2^ units. While Kernel PCA test *R*^2^ values are not trivial in absolute terms—reaching 0.67 at *k* = 100—they fall well below the linear baseline under identical experimental conditions. This result suggests that for donorheld-out generalization in this dataset, the linear assumption is not a limitation but may in fact be an advantage: the structured, low-dimensional geometry captured by PCA appears to generalize more robustly across patients than the nonlinear RBF manifold, which may overfit donor-specific curvature in the training set.

We additionally note that the linear PCA results also provide indirect evidence on this point: under strict donor-held-out evaluation, FBG achieves test *R*^2^ of 0.83–0.97 across RCC and AD. A model that fails to capture meaningful transcriptomic structure would not generalize at this level to completely unseen donors.

##### Submodularity Diagnostic Under Kernel PCA

We repeated the submodularity diagnostic (2,000 randomized triplet tests per seed, five seeds per dataset) replacing linear PCA with RBF Kernel PCA across four of the five datasets analyzed in this paper. For the fifth dataset (GSE227734), the Kernel PCA eigendecomposition failed due to its highly restricted sample size (only 338 cells, compared to 13,000–200,000+ cells in the other cohorts), which causes severe numerical instability when extracting 50 components. Across the four evaluable datasets, mean violation rates under Kernel PCA were: 14.12% (GSE138852), 3.81% (GSE314072), 9.83% (GSM8652069), and 0.02% (GSE308624). In all cases, the mean marginal difference Δ(*g* | *S*^*′*^) − Δ(*g* | *S*) remained negative, and the near-submodular verdict was confirmed. These results demonstrate that diminishing returns in the ridge reconstruction objective persist under nonlinear embeddings, and are not an artifact of the linear PCA assumption.

#### A.8 Empirical Estimation of the Sub-modularity Ratio via Restricted Gram Matrix Conditioning

The theoretical connection between greedy approximation guarantees and the submodularity ratio *γ*_*U,k*_ is established by Theorem 1 of Elenberg et al. [2018], which provides the bound *γ*_*U,k*_ *≥ m*_|*U*|+*k*_*/M*_|*U*|+*k*_, where *m* and *M* denote the restricted strong convexity (RSC) and restricted strong smoothness (RSM) parameters of the objective over sparse supports. Computing exact RSC/RSM constants over all *k*-sparse subsets is NP-hard in general; we therefore adopt an empirical estimation approach.

For our ridge-regularized objective, the Hessian restricted to a support *S* is 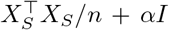, whose minimum and maximum eigenvalues directly correspond to the RSC parameter *m* and RSM parameter *M*, respectively. We estimate both quantities by computing the minimum and maximum eigenvalues of 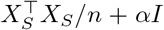 over 50 randomly sampled sparse gene subsets per seed, for all sparsity levels *k* ∈ {10, 25, 50, 100}, across five random donor splits and both the RCC (GSE314072) and AD (GSE138852) datasets (*α* = 1.0, matching the main experimental configuration). While this does not constitute a formal worst-case lower bound, the consistency of the resulting ratio across 250 independent random supports per condition suggests the objective is well-conditioned over typical sparse supports encountered in practice—which is the operationally relevant regime for greedy selection.

The empirical RSC/RSM ratio, providing an approximation of *γ* by Theorem 1 of Elenberg et al. [2018], ranged from 0.84 to 0.30 across *k* ∈ {10, 25, 50, 100} for RCC, and 0.85 to 0.36 for AD. The ratio decreases with *k* as expected, since larger sparse supports admit greater spectral spread in the Gram matrix. Critically, the ratio remains bounded away from zero at all evaluated budgets, confirming that the theoretical greedy approximation guarantee *f* (*S*_greedy_) *≥* (1 − *e*^−*γ*^)*f* (*S*^*∗*^) holds with non-trivial *γ* throughout the sparsity levels evaluated in this work [Das and Kempe, 2011, Bian et al., 2017].

